# Basal Ganglia role in learning rewarded actions and executing previously learned choices: healthy and diseased states

**DOI:** 10.1101/616854

**Authors:** Garrett Mulcahy, Brady Atwood, Alexey Kuznetsov

**Affiliations:** Department of Mathematics, Purdue University; Departments of Psychiatry and Pharmacology & Toxicology, IUSM; Department of Mathematical Sciences, IUPUI and Indiana Alcohol Research Center, IUSM

## Abstract

The basal ganglia (BG) is a collection of nuclei located deep beneath the cerebral cortex that is involved in learning and selection of rewarded actions. Here, we analyzed BG mechanisms that enable these functions. We implemented a rate model of a BG-thalamo-cortical loop and simulated its performance in a standard action selection task. We have shown that potentiation of corticostriatal synapses enables learning of a rewarded option. However, these synapses became redundant later as direct connections between prefrontal and premotor cortices (PFC-PMC) were potentiated by Hebbian learning. After we switched the reward to the previously unrewarded option (reversal), the BG was again responsible for switching to the new option. Due to the potentiated direct cortical connections, the system was biased to the previously rewarded choice, and establishing the new choice required a greater number of trials. Guided by physiological research, we then modified our model to reproduce pathological states of mild Parkinson’s and Huntington’s diseases. We found that in the Parkinsonian state PMC activity levels become extremely variable, which is caused by oscillations arising in the BG-thalamo-cortical loop. The model reproduced severe impairment of learning and predicted that this is caused by these oscillations as well as a reduced reward prediction signal. In the Huntington state, the potentiation of the PFC-PMC connections produced better learning, but altered BG output disrupted expression of the rewarded choices. This resulted in random switching between rewarded and unrewarded choices resembling an exploratory phase that never ended. Our results reconcile the apparent contradiction between the critical involvement of the BG in execution of previously learned actions and yet no impairment of these actions after BG output is ablated by lesions or deep brain stimulation. We predict that the cortico-BG-thalamo-cortical loop conforms to previously learned choice in healthy conditions, but impedes those choices in disease states.

**Author summary:** Learning and selection of a rewarded action, as well as avoiding punishments, are known to involve interaction of cortical and subcortical structures in the brain. The subcortical structure that is included in this interaction is called Basal Ganglia (BG). Accordingly, diseases that damage BG, such as Parkinson and Huntington, disrupt action selection functions. A long-standing puzzle is that abolition of the BG output that disconnects the BG-cortical interaction does not disrupt execution of previously learned actions. This is the principle that is suggested to underlie standard Parkinsonian treatments, such as deep brain stimulation. We model the BG-cortical interaction and reconcile this apparent contradiction. Our simulations show that, while BG is necessary for learning of new rewarded choices, it is not necessary for the expression of previously learned actions. Our model predicts that the BG conforms to previously learned choice in healthy conditions, but impedes those choices in disease states.

## Introduction

The basal ganglia (BG) is a complex network of excitatory and inhibitory neurons located in the deep brain of most vertebrates that controls action selection (see e.g. (1)). The BG is comprised of the dorsal striatum, external and internal portions of the globus pallidus (GPe, GPi), subthalamic nucleus (STN) and substantia nigra (2). It is traditionally implicated in motor control since BG lesions are associated with movement disorders (3–5). The BG is a shared processing center involved in a broad spectrum of motor and cognitive control (2,6). A cortico-BG-thalamo-cortical neurocircuit loop is suggested to be the structure that provides this control (7,8). However, understanding how this loop functions remains far from complete and requires more experimental and theoretical studies.

The BG is also widely recognized for its involvement in learning (9–11). Reinforcement learning is recognized as the mechanism that establishes behavioral responses for rewards, such as food or drugs of abuse and is altered in numerous disorders and disease states including Parkinson’s disease (12–14). Reinforcement learning is based on communication between midbrain dopamine neurons and the striatum (15,13), specifically ventral tegmental area projections to ventral striatum and substantia nigra pars compacta (SNc) projections to dorsal striatum (16–18). Dopamine (DA) released by dopaminergic VTA and SNc inputs to striatum signals the difference between received and expected rewards – the reward prediction error (RPE) (14). RPE encoding in VTA-NAc neurocircuits involves prediction of reward value which in turn feeds back to both VTA and SNc dopamine neurons (15). Given its role in motor control, the SNc-dorsal striatum component of the BG translates RPE into action: the hypothesized critic-actor roles of these two dopaminergic projections (15,14,19,20). If the RPE is positive, additional DA release leads to positive reinforcement of the preceding action; if the error is negative (expected more than received), a pause in DA release leads to negative reinforcement and blocks the action. As a mechanism for this control, DA modulates plasticity of synaptic projections from the cortex to striatal medium spiny neurons (MSNs) (21,22). As a reflection of the bidirectional DA modulation, there are two types of MSNs. Those that are responsible for promoting movement are part of the BG direct pathway and express D1-type dopamine receptors (GO, D1-MSNs) and those that inhibit movement are part of the BG indirect pathway and express D2 dopamine receptors (NO-GO, D2-MSNs) (23–25). Indirect and direct BG pathways respectively inhibit or disinhibit the thalamocortical relay neurons responsible for producing particular movements (26–28). The coordination of activity within the two types of MSNs determines action (29–31). Within the BG loops, synaptic plasticity of corticostriatal projections is a key node in the learning of rewarded choices (9–11,22).

The BG is suggested to remain involved in action selection after the action-reward association is learned and control the transition from goal-directed to habitual choices (8,32). On the other hand, clinical interventions for Parkinson disease (PD) do not cause impairments in goal-directed or habitual movements (33–35). Specifically, GPi lesions and deep brain stimulation, which disrupt the main output of the BG, are used to improve motor functions. This observation gave rise to a hypothesis that the BG play a critical role in learning, but not in the expression of already learned actions or choices (36,37). These choices are suggested to instead be stored in synaptic connections within cortex. This hypothesis apparently contradicts the suggested involvement of the BG in executing actions learned previously. Therefore, it is essential to fill in this knowledge gap by further investigating the role of the BG in goal-directed and habitual choices.

This paper presents a computational model of the cortico-BG-thalamo-cortical loop involved in a two-choice instrumental conditioning task (32). This task is standard for assessing action-reward association in animals and humans. Our model design is similar to a previously published design (37,38), but focused on choice selection. We implemented two synaptic mechanisms that can mediate learning: reward-related plasticity of corticostriatal synapses (39) and activity-dependent Hebbian plasticity (40,41) of cortico-cortical synapses. To elucidate the role of the BG in Parkinson’s and Huntington diseases, we calibrate the model to reflect the altered BG connectivity documented for these diseases and simulate these changes in BG activity.

## Results

We simulated the same standard two-choice IC and reversal task in three conditions: Healthy, Parkinsonian, and Huntington’s BG. Fig. 1 presents a schematic diagram of nuclei and connections within the BG and their connections with cortices. The model is described in detail in Materials and Methods. The models received a stimulus (CS) that activates prefrontal cortical (PFC) neurons for all 500 trials. We say that the network chooses action 1 if the premotor cortical (PMC) neural group 1 displays greater activity than the PMC group 2. For reversal training, after action 1 is rewarded in trials 1 through 199, for trials 200 through 500, action 2 was rewarded instead. We analyze and compare the learning and reversal performance in the three model states below.

**Figure 1:**
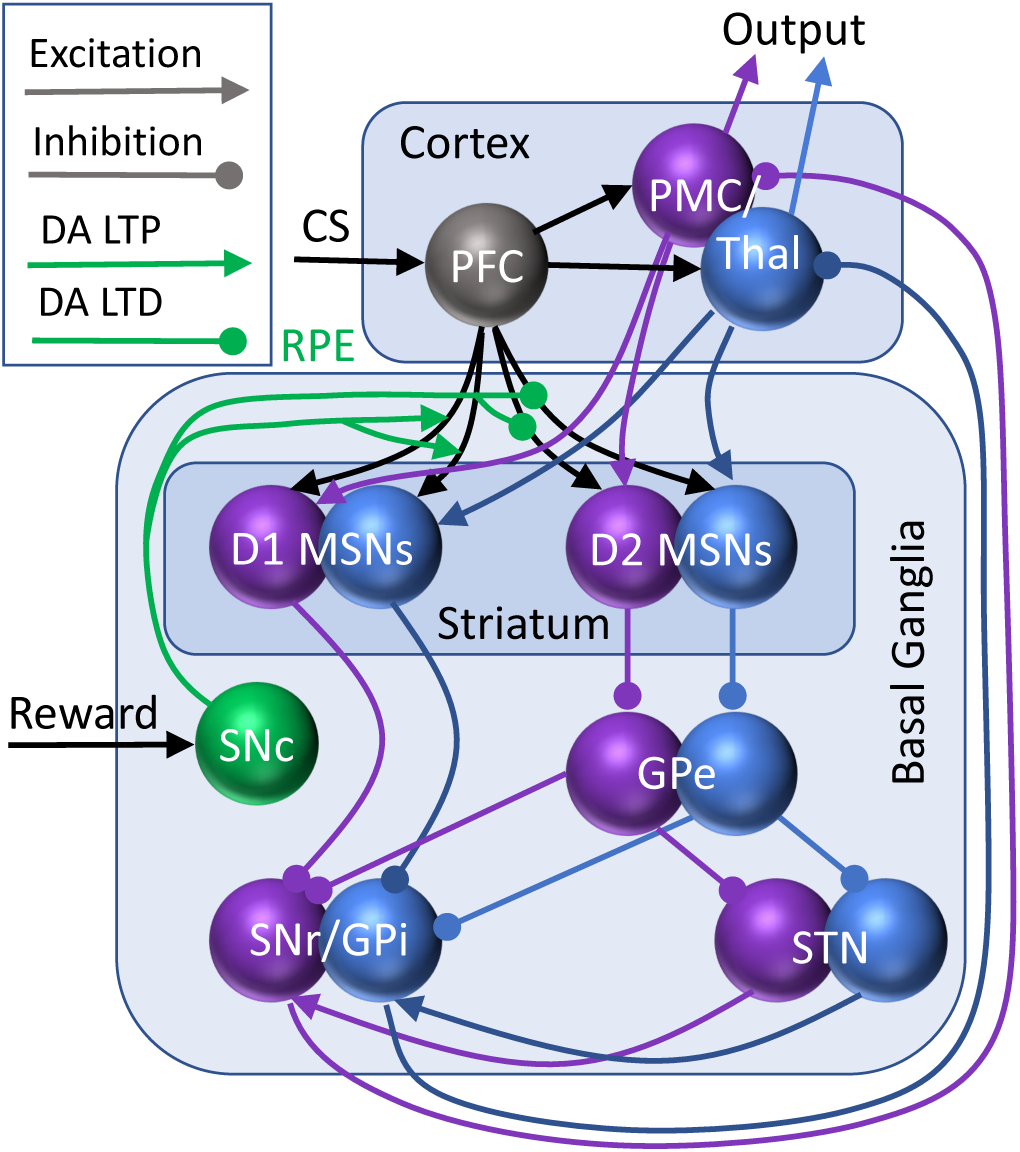
The structure of the cortico-basal ganglia-thalamo-cortical loop model. The BG receives inputs from the prefrontal cortex (PFC) signaling the conditioning stimulus (CS) as well as reward inputs via substantia nigra pars compacta (SNc). The SNc forms a dopamine reward prediction error (RPE) signal, which governs plasticity of the connections from the PFC (DA LTP/LTD; green). The BG input structure, striatum, contains medium spiny neurons (MSNs), which cluster in 2 subtypes: D1 and D2 dopamine receptor-containing (direct and indirect pathways respectively). The rest of the nuclei are the globus pallidus external (GPe), subthalamic nucleus (STN), and the output structures: substantia nigra pars reticulata and globus pallidus internal (SNr/GPi). The loop is completed by connections from and to premotor cortices/thalamus (PMC/Thal). The two channels of the loop are colored purple/blue.

### Healthy BG facilitates learning of rewarded choices

Fig. 2A shows choices made in the simulations: a higher activity of the PMC1 manifests choice 1 and vice versa. The graph shows the activity at the end of each trial, which is taken to be 750 msec long. On early trials, the choice is made randomly due to random initial conditions in the PMC network and mutual inhibition of PMC1 and PMC2. This reproduces the exploration phase, where the information about reward is collected (42,43). The modeled animal receives an unexpected reward every time it chooses action 1 (PMC1 on top). Within 20 trials, the system starts to consistently choose the rewarded action, and only a few exploratory deviations are made after that. On trial 200, we switch the simulated task to reversal: action 2 is rewarded instead. This quickly leads to reestablished exploratory behavior, and then locks the system to the rewarded choice, with occasional exploratory returns to choice 1. As explained below, our model allows for detailed analysis of the mechanism of this learning.

**Figure 2:**
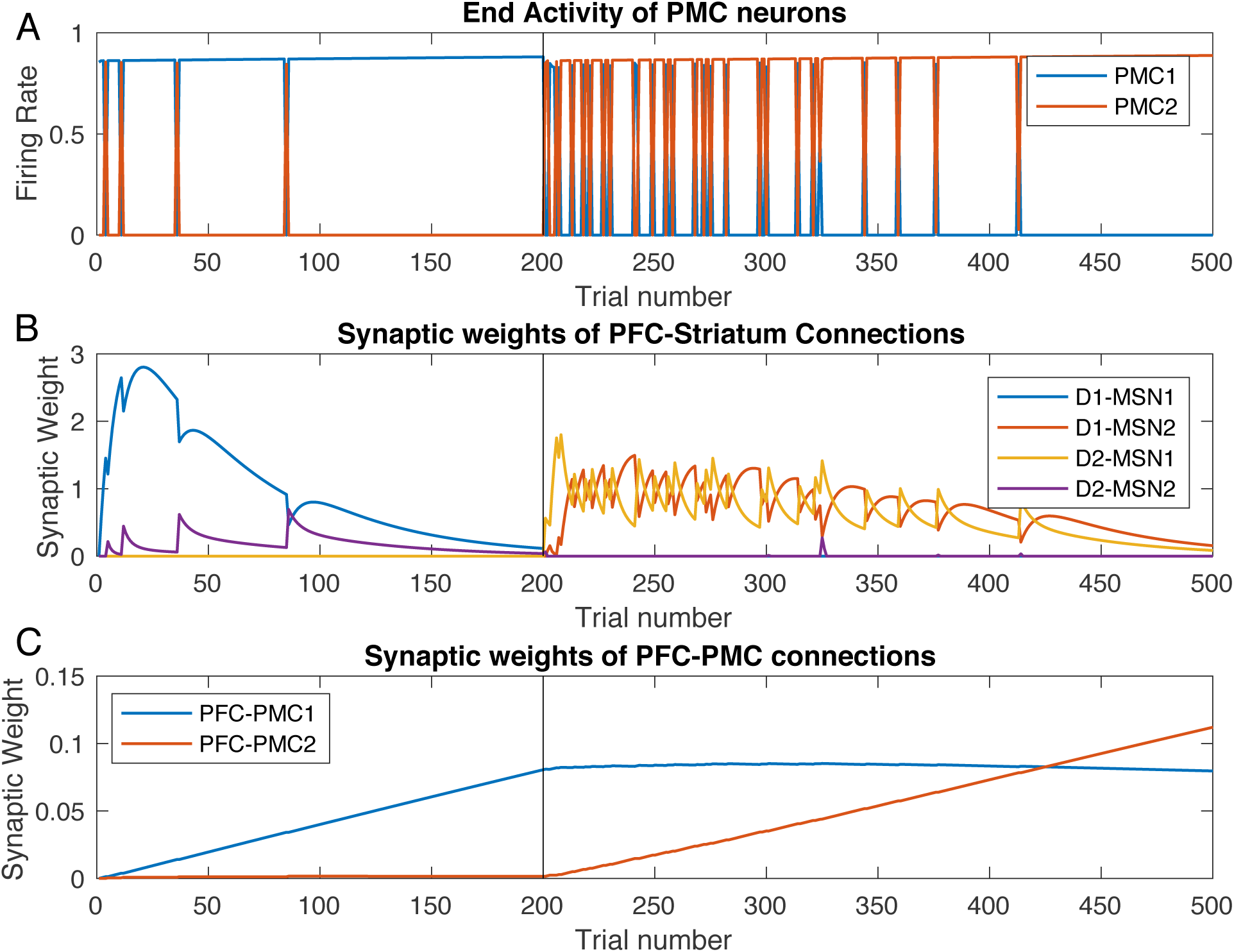
Healthy BG facilitates learning of the initial task and reversal. Trial-by-trial dynamics of the PFC activity and underlying modulation of synaptic weights in the Healthy BG model. Trials 1-199:initial learning; trials 200-500: reversal (A) A higher activity of PMC1 (blue) manifests choice 1, whereas higher activity of PMC2 manifests choice 2. (B) Synaptic weights of the PFC to striatum connections. (C) Synaptic weights of the PFC to PMC connections.

Two mechanisms facilitate learning of the rewarded choice – one fast and one slow. The first mechanism is the potentiation of the PFC-to-striatum synaptic connections (Fig. 2B). The unexpected reward creates a positive RPE encoded by SNc DA signaling and potentiates PFC connections to all D1-MSNs (Fig. 2B). Note that the connections to D2-MSNs are potentiated much later (see below). Importantly, whereas the DA signal itself is not selective for MSNs specific to the rewarded action, DA-mediated potentiation of PFC-MSN synapses is selective. What makes potentiation selective is the level of activation of the corresponding MSN: in the initial trials the reward is granted only if choice 1 is selected, that is when PMC1 activity is greater, and this happens only when the corresponding D1-MSN is active (due to static synaptic connections from PMC to MSNs specific for each choice). Since synaptic plasticity explicitly depends on the activity of the postsynaptic neuron, PFC-to-MSN1 connections are potentiated much more strongly than other MSN connections (Fig. 2B). This further selectively activates D1-MSN1s. Thus, excitation of D1-MSNs neurons associated with choice 1 increases due to direct excitation from the PFC associated with the stimulus. The increased activity level of D1-MSN1s inhibits downstream GPi1 neurons and, consequently, disinhibits the PMC1 neural group (Fig. 3).

**Figure 3:**
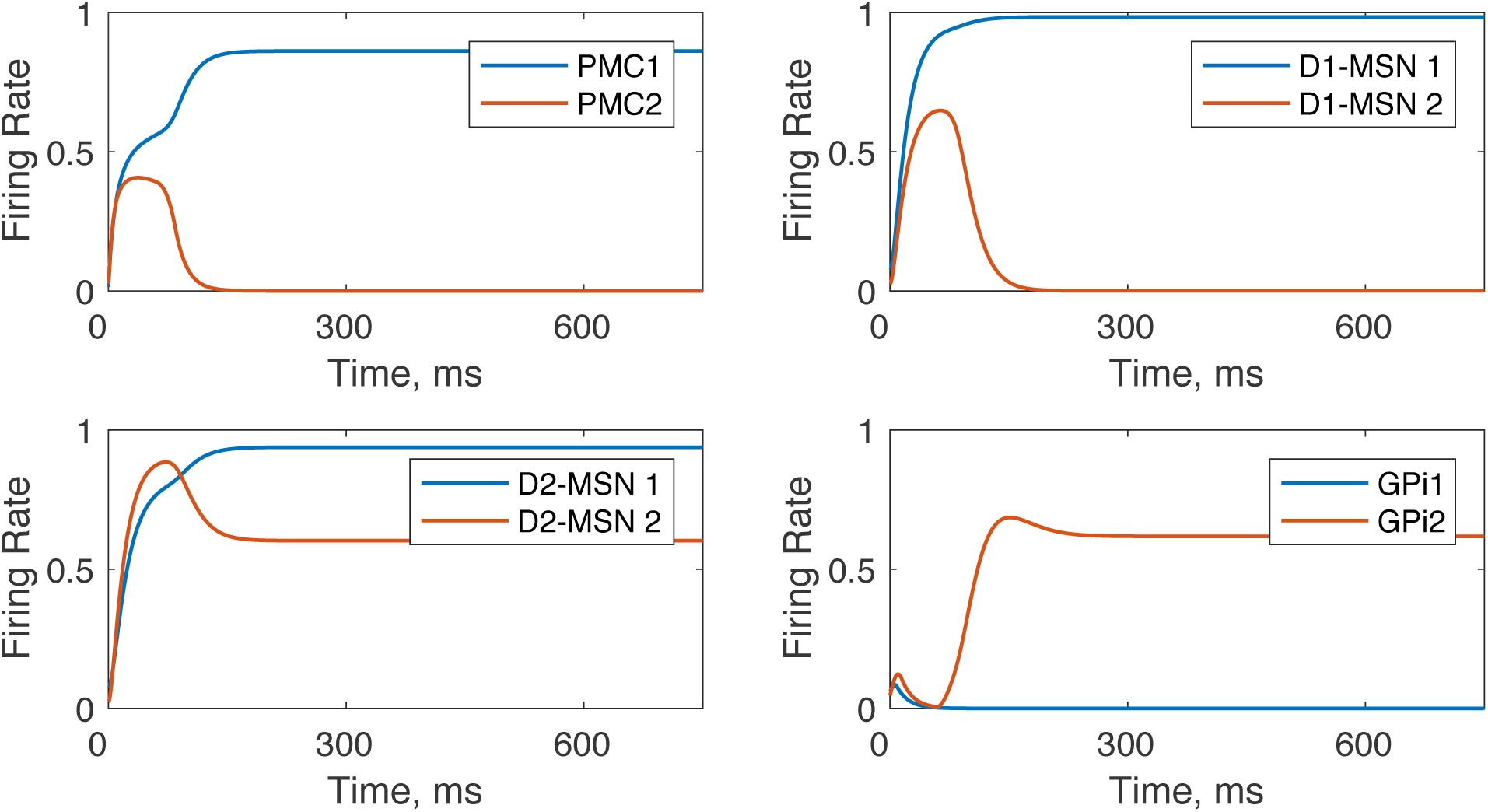
Within-trial dynamics of neural activity in the model with healthy BG. The network is biased towards option 1 as the PFC-D1-MSN1 and PFC-D2MSN2 connection weights are both set at 0.7, which corresponds to a trial in late initial learning phase (∼100). Activation of the D1-MSN1 group inhibits GPi1 neurons, and thus disinhibits PMC1. GPi2 neurons remain excited and inhibit PMC2.

The PFC to D2-MSN connections are potentiated much later in the process (Fig. 2B, purple) and further reinforce the activity of PMC1. The potentiation delay is because a negative RPE is required for activation of the D2 MSNs, which is formed after the expected reward builds up and a nonrewarded action is selected by chance. Then, every choice that is not followed by the expected reward activates the corresponding indirect pathway (i.e. D2-MSN2), which excites the downstream GPi2 neurons, and consequently inhibits the PMC2 activity (Fig. 3). This blocks the nonrewarded action and helps to lock the choice to the rewarded action. Co-activation of the two mechanisms is sufficient to lock the choice to the rewarded action.

During subsequent repetitions of the same trial, the PFC-MSN connection strength starts to decrease and approaches zero (Fig. 2B trials 80 to 200). However, the persistence of the rewarded choice remains intact (Fig. 2A). The mechanism for this is the growth of direct PFC-PMC1 connections (Fig. 2C) via classical reward-independent Hebbian synaptic plasticity: the two neural groups are co-active most of the time. This transition from PFC-MSN to PFC-PMC connections as a supporting mechanism for the rewarded choice occurs after the number of repetitions is in the order of a hundred (Fig. 2). Therefore, the model shows that direct cortico-cortical connections are responsible for the choice of the rewarded action after long training.

We next analyzed the behavior of the model when we began rewarding a choice different from the choice the model had been previously conditioned to make; this learning task is called reversal learning (44). Beginning at trial 200, we rewarded the model for selecting the other action (choice 2). Thus, starting at trial 200 the model mimics omission of a reward, which acts as an unexpected punishment (negative reward) for selecting action 1. This punishment potentiates synaptic connections from the PFC to D2-MSNs associated with action 1 (D2-MSN1, Fig. 2B yellow), and, slightly later, to D1-MSNs associated with action 2 (D1-MSN2, Fig 2B red). This engagement of both direct and indirect pathways offsets the model bias for action 1 and quickly sends the model into another exploratory phase. As Fig. 2A demonstrates, between trials 200 and 300 the model is randomly choosing between the two actions. It is important to note that, in accordance with others’ findings (45,46), this second exploratory phase lasts longer than the initial exploratory phase. During reversal, the new potentiation of PFC-MSN connections is not enough to effectively overcome the bias for the initially learned choice and ensure choosing the newly rewarded option. The reversal exploratory phase ends only when the PFC-PMC2 connections become as strong as PFC-PMC1 and remove the bias (Fig. 2). Thus, the longer exploratory phase during reversal occurs because the model must first overcome its bias for the previously learned choice and then develop a new stimulus-choice 2 association.

The reversal mechanism relies more on the D2-MSN, indirect pathway and less on the D1-MSN, direct pathway than the initial learning. Due to the potentiated PFC-PMC1 connection, the system continues choosing option 1, even though it’s not rewarded. This generates a negative reward prediction error (Fig. 4) and potentiates PFC connections to the D2-type neurons associated with action 1 (D2-MSN1; Fig. 2B yellow). The connection of PFC to D1-MSN2, which conducts the GO signal for the choice 2 lags by several trials (Fig. 2B, red), during which the exploratory phase begins and allows finding the new rewarded option. The connections to the D1 MSNs do not potentiate as strong during reversal as those during initial learning. Their temporal profile closely matches the positive RPE signal, which also stays significantly lower during reversal compared to the initial learning (Fig. 2B blue and red, Fig. 4 RPE). As a result, the reversal learning engages direct and indirect pathways at a comparable strength (Fig. 2B, yellow and red), whereas during initial learning the direct pathway is engaged much stronger than the indirect (Fig. 2B, blue and purple).

**Figure 4:**
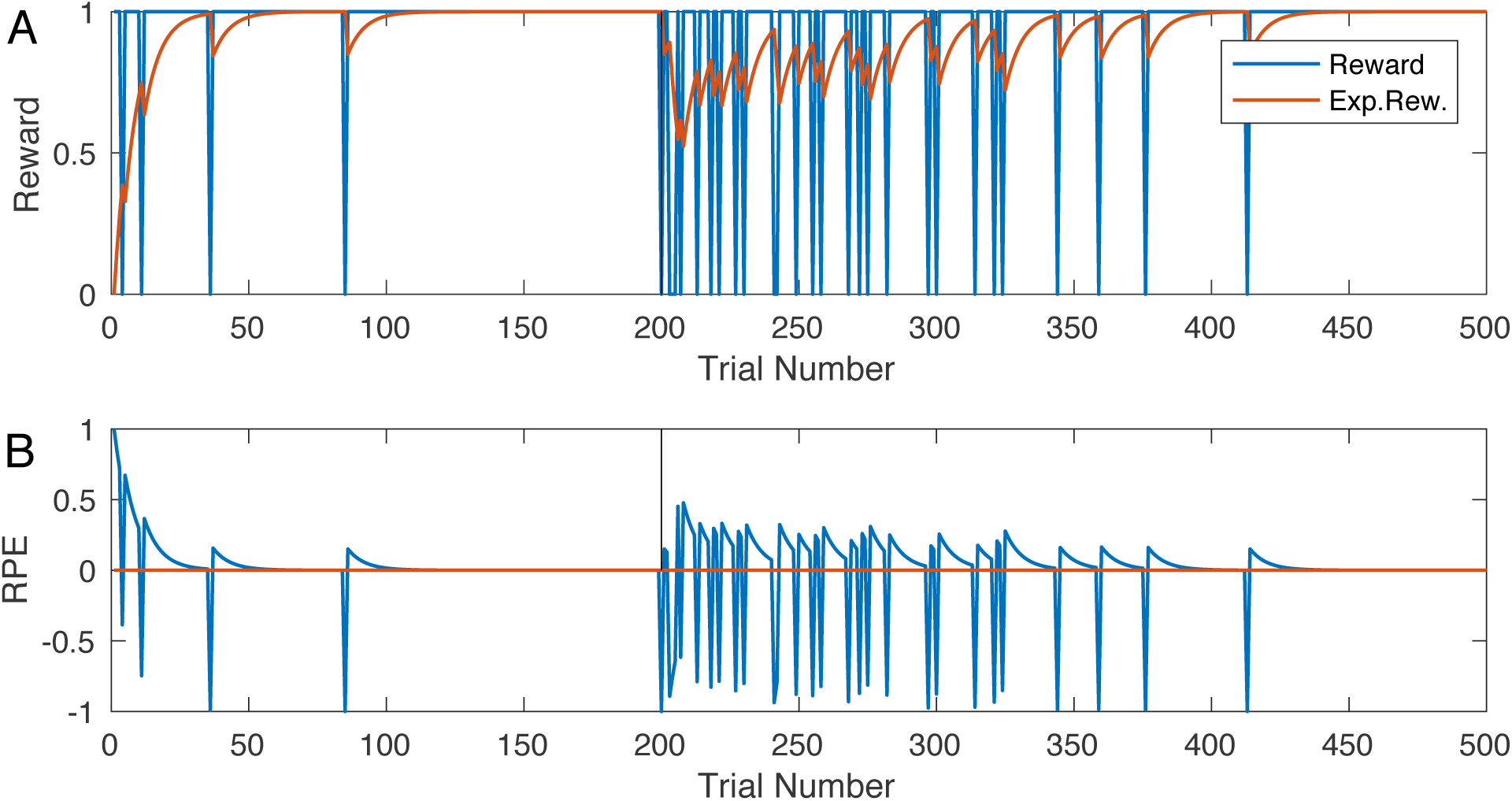
Reward, expected reward (A), and the RPE (B) during initial learning and reversal trials in the model with healthy BG. As before, reversal starts at trial 200 (vertical black line). Note a greater RPE at the beginning of the initial learning compared to the reversal.

### Mild Parkinsonian BG: impeded learning and spontaneous oscillations

Our simulations (Fig. 5) show drastic difference in dynamics of the PMC neurons during initial learning and reversal in the model with mid-parkinsonian BG. During both phases, learning is severely impaired. First, the choice remains random for approximately the first 120 trials. Second, the model does not reliably choose the rewarded option even after this period, although the rewarded option is chosen on a much greater number of trials (Fig. 5A blue above red in the initial learning and vice versa in the reversal). Third, the activity of the PMC neurons is overall reduced compared to that in the model with healthy BG, and the trial-to-trial variations of this activity are drastically increased, even when only trials with the same choice are considered.

**Figure 5:**
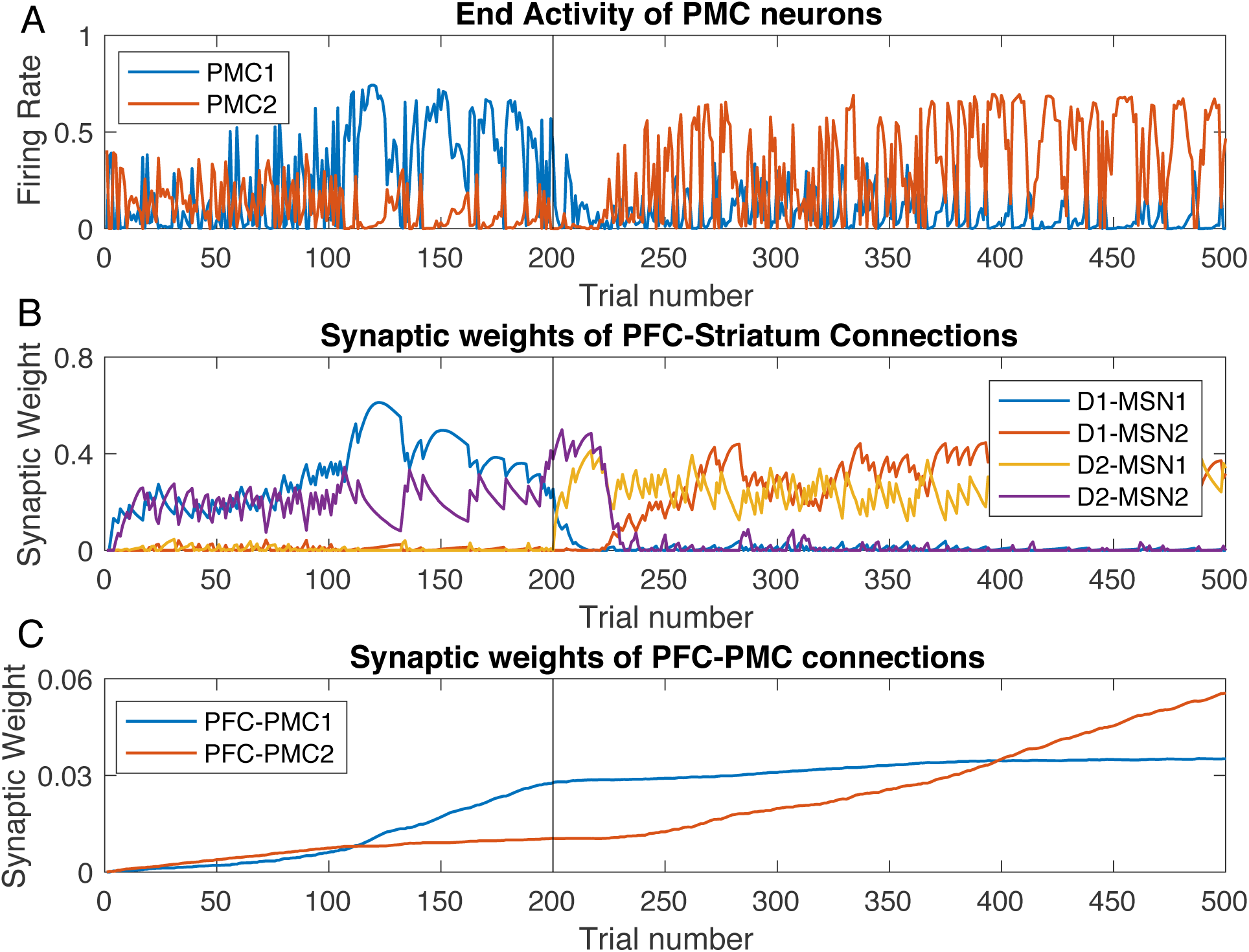
Decreased learning performance and increased variability of PMC activity in the model with mild-parkinsonian BG. Trial-by-trial dynamics of PMC activity (A) and underlying modulation of synaptic weights (B,C) in the model with mild-parkinsonian BG state. Notation is the same as in Fig. 2. Note the difference in scale in panels (B) and (C) compared to Fig. 2

The underlying dynamic of the synaptic weights is also significantly altered. During initial learning, both direct pathway for choice 1 and indirect pathway for choice 2 are activated at a similar level (Fig. 5B), and this level is much lower than in the model with healthy BG (Fig. 2B). The latter follows directly from the reduced SNC signaling (by 70%), which decreases the RPE and, thus, impedes potentiation of PFC-MSN connections. Since both PMC neural groups are active at a similar level, both connections from PFC are potentiated (Fig. 5C), and the system does not develop a preference for the rewarded choice. After trial 80, the rewarded choice starts to prevail as the PFC-PMC connections reflect the preference for choice 1. However, the PFC-PMC1 connection does not achieve the level reached in the model with healthy BG (Fig. 2C) within the 200 trials designated for initial learning. Hence, exploration between the choices persists for all 200 trials, and the prevalence of the rewarded choice requires the persistent activation of PFC-MSN connections. Therefore, the model with mild parkinsonian BG is capable of learning the choices, but the effective learning rate is much lower.

The low levels of PFC-PMC connections persist into the reversal phase too and never reach the levels shown by the model with healthy BG even though plasticity rules of the PFC-PMC connections remain the same in both models. Therefore, our modeling predicts that the mild-parkinsonian BG does not allow for the proper potentiation of the PFC-PMC connections, and this leads to impaired learning. Interestingly, the reversal phase starts with activation of both indirect pathways simultaneously (Fig. 5B, purple and yellow). This suppresses the activity of both PMC neural groups, blocks any choice and blocks changes in the PFC-PMC synaptic weights. Only after some 40 trials, the NO-GO signal for choice 2 is replaced by a GO (Fig. 5 purple and red). Thus, the model with mild-parkinsonian BG predicts that the exploratory phase at the beginning of the reversal learning is replaced by blockade of any choice, and this further impedes learning.

Perhaps the most interesting change in the model with parkinsonian BG is the drastic increase in the trial-to-trial variability of the PMC neurons (Fig. 5A). To explain the mechanism of this variability, we considered within-trial dynamics of activity for all neural groups in the model. Fig. 6 shows these dynamics for the PMC neurons and MSNs in the healthy vs. parkinsonian BG models. In the healthy case activity levels come to an equilibrium, while in the parkinsonian case, they engage in persistent oscillations. The anti-phase for the oscillations in the neural groups corresponding to the choice 1 and 2 is due to mutual competition (inhibition) between PMC1 and PMC2 groups. The oscillations arise from the negative feedback loop that the BG, and in particular its indirect pathway, provides for the activity of each PMC neural group. Indeed, the static PMC to D2 MSN connections, which constitute this negative feedback, are stronger in the parkinsonian case (*W_PMC–D_*_2_, in Table 2). The period of these oscillations is approximately 210 ms, which is 4.7 Hz. No potentiation in the PFC-PMC and PFC-MSN connections within the ranges in Fig. 5 B and C suppress the oscillations (data not shown). Therefore, the simulations predict that the trial-to-trial variability of the PMC neurons in the model with parkinsonian BG is caused by robust within-trial oscillations in the activity of all neuron groups in the model.

**Figure 6:**
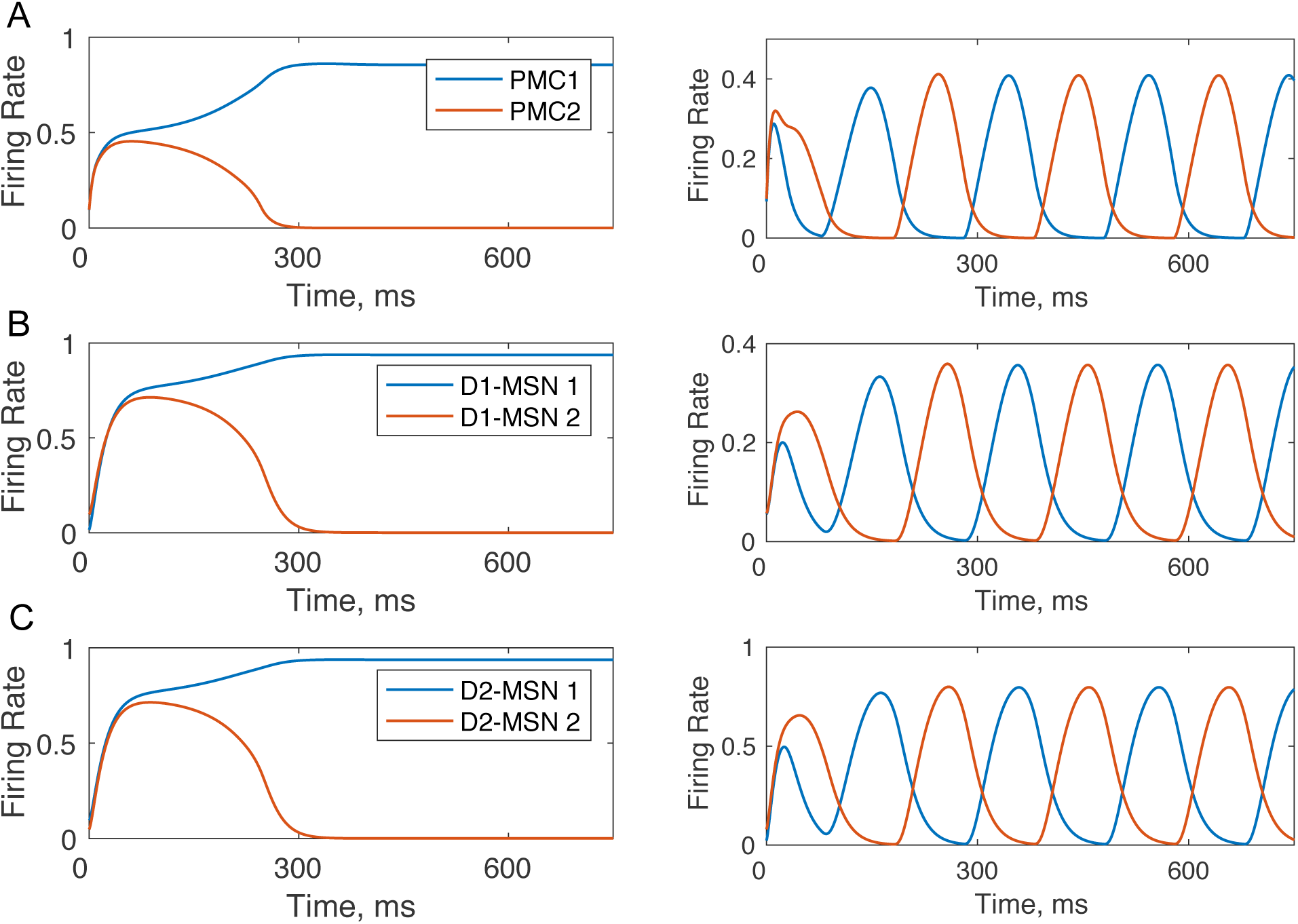
Within-trial dynamics of neural activity in the model with healthy (left) and parkinsonian (right) BG. Panels A, B, and C show firing rates for PMC,D1 MSNs and D2 MSNs respectively. In the healthy case, the firing rates equilibrate within 500 ms. In the parkinsonian case, oscillations in the firing rate emerge and persist. All plastic synaptic connections are set to zero to simulate the state of no bios towards any choice.

**Table 1:** Parameters of the healthy BG model state

| Parameter | Value used in this model |
| --- | --- |
| $input\_pfc$ | 3.0 |
| $w_{PFC-D1_m}$ & $w_{PFC-D2_m}$ | Randomly set between 0 and 0.001, updated after each trial |
| $w_{PMC-D1}$ | 2.0 |
| $w_{PMC-D2}$ | 2.0 |
| $dr_{GPe}$ | 2.0 |
| $w_{D2-GPe}$ | 2.0 |
| $dr_{STN}$ | 1.0 |
| $w_{GPe-STN}$ | 1.0 |
| $dr_{GPi}$ | 0.2 |
| $w_{D1-GPi}$ | 1.4 |
| $w_{STN-GPi}$ | 1.6 |
| $dr_{PMC}$ | 1.3 |
| $w_{PFC-PMC_m}$ | Initial 0; varies with trials |
| $w_{GPI-PMC}$ | 1.8 |
| $w_{PMC_m-PMC_n}$ | 1.6 |
| $\lambda$ | 0.0005 |
| $\lambda_{CM}$ | 0.0005 |

**Table 2:** Changes in the parameters of the model that reproduce Parkinsoninan BG state.

| Parameter | Value in Healthy state | Value in mild Parkinsonian state | Justifying literature |
| --- | --- | --- | --- |
| $w_{PMC-D1}$ | 2.0 | 1.0 | (65,66) |
| $w_{PMC-D2}$ | 2.0 | 3.0 | (65,66) |
| $dr_{STN}$ | 1.0 | 1.1 | (71,72) |
| $dr_{GPI}$ | 0.2 | 0.3 | (73,67,74) |
| $w_{D1-GPI}$ | 1.4 | 1.0 | (73,67,74) |
| $w_{STN-GPI}$ | 1.6 | 2.0 | (73,67,74) |

In order to model the impact of BG DBS or surgical interventions on performance and learning in PD, we performed additional simulations of the PD model in which the BG signal to PMC was ablated from trial 150 till the end (Fig. 7). In this period, the variability of the PMC activity vanishes completely. Furthermore, the PFC-striatal connections no longer exert any influence on the choices, but the PFC-PMC connections are strong enough to lock the choice to the rewarded option, and the cortical connections increase further at a greater rate. After the reversal on trial 200, however, the changed values of the choices remain unnoticed by the system, the choice remains locked on the now unrewarded option, and the cortical connections supporting this choice keep rising. In this state, behavior improves, but learning is impaired.

**Figure 7:**
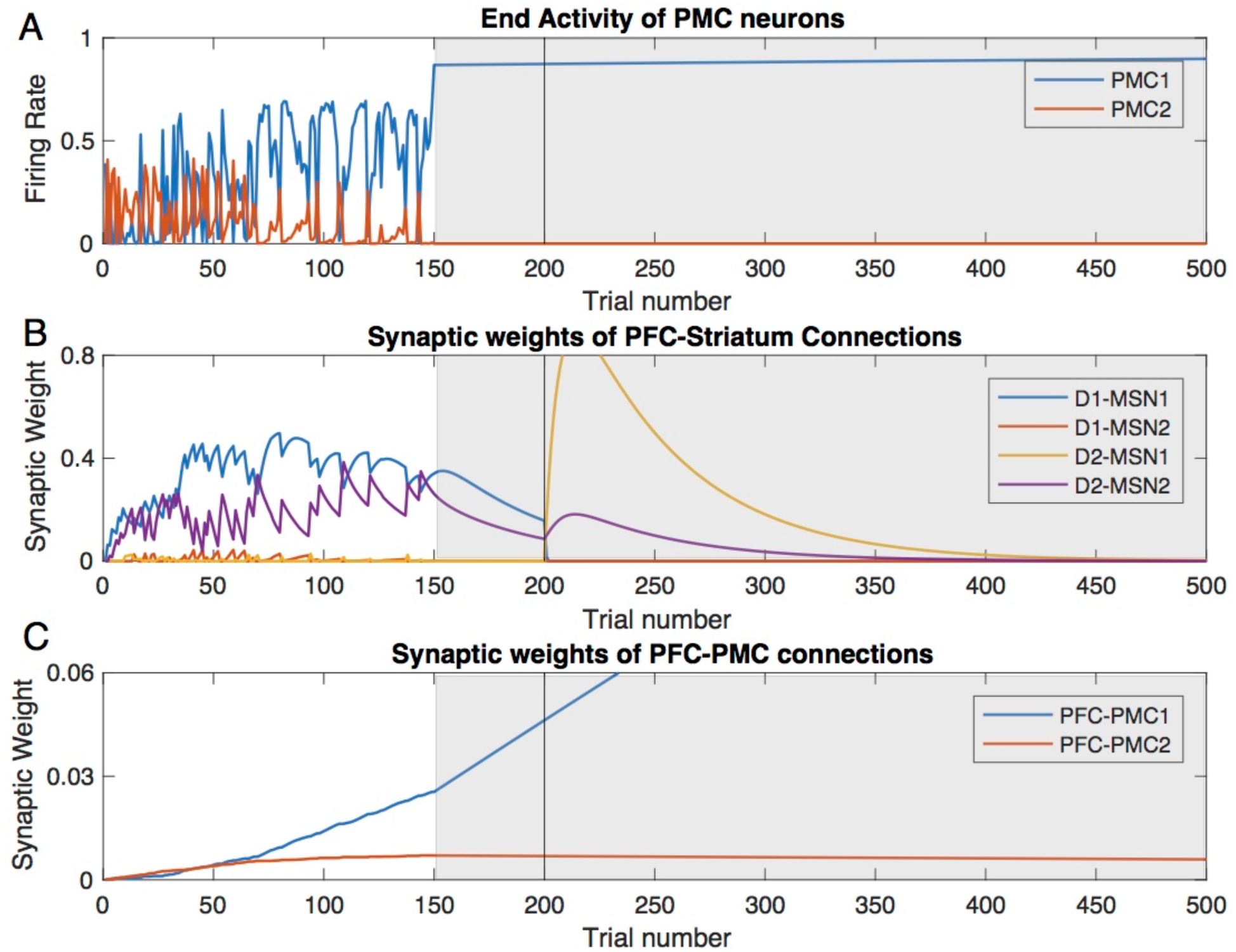
In the PD state model, the variability of PMC activity and switching between choice 1 and 2 cease at the DBS onset. Trial-by-trial dynamics of the PMC activity and underlying modulation of synaptic weights in the PD BG model with simulated DBS starting at trial 150. Same notation as in Fig. 2. (A) The levels of PMC1 and PMC2 activity (choice 1 vs. 2) at the end of each trial (B) Synaptic weights of the PFC to striatum connections reflect rewarded choices. (C) Synaptic weight of the PFC to PMC1 connection keep growing after DBS onset, and during reversal.

### Grade 2 Huntington’s Disease BG state: persistent exploratory behavior

If the above case of Parkinson’s disease is associated with strengthening the indirect pathway, in the case of Huntington’s disease the connections in the indirect pathway become weaker (Table 3). The major difference with the healthy BG model is that the trial-to-trial dynamics of the PMC neural groups looks like the exploratory phase never ends (Fig. 8A). At the same time, we see from the synaptic weights (Fig. 8B and C) that choice-reward contingencies are learned almost as effectively as in the healthy case (Fig. 2), although the synaptic weights are somewhat lower. The differences are the activation of the indirect pathway for choice 2 lingering at the beginning of the reversal phase (Fig. 8B purple) and the persistence of the PFC-MSN connections similar to the parkinsonian case. The latter, however, is not a cause but rather a consequences of the continuous exploratory choices that bring no reward. Therefore, despite the efficacious learning (Fig. 8C), choice behavior is impaired relative to control (Fig. 8A).

**Figure 8:**
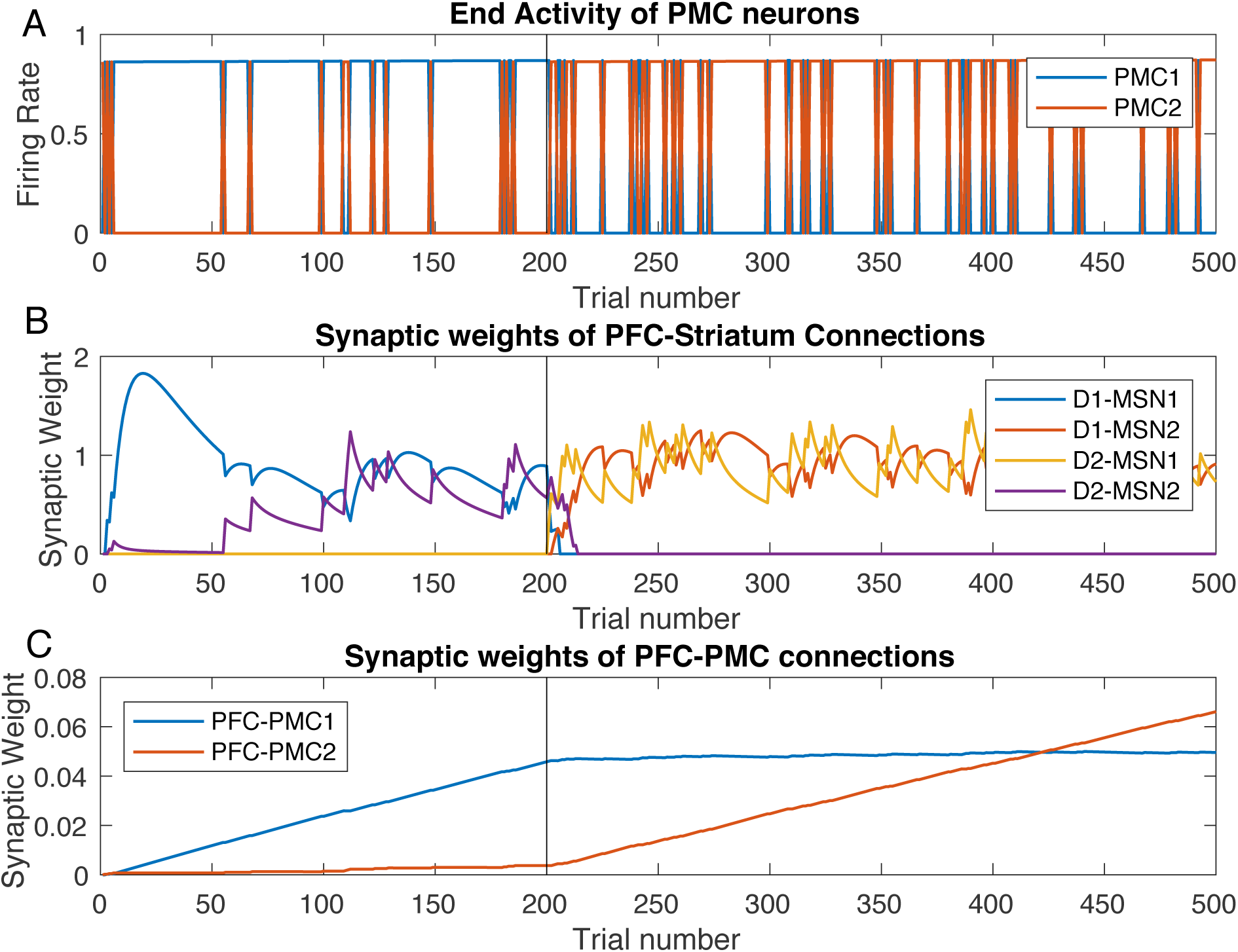
Random switches between rewarded and unrewarded options persist in the model with Huntington state BG. Trial-to-trial dynamics of PFC neural activity (A) and underlying dynamics of synaptic weights (B,C). The notation is the same as in Fig. 2.

**Table 3:**
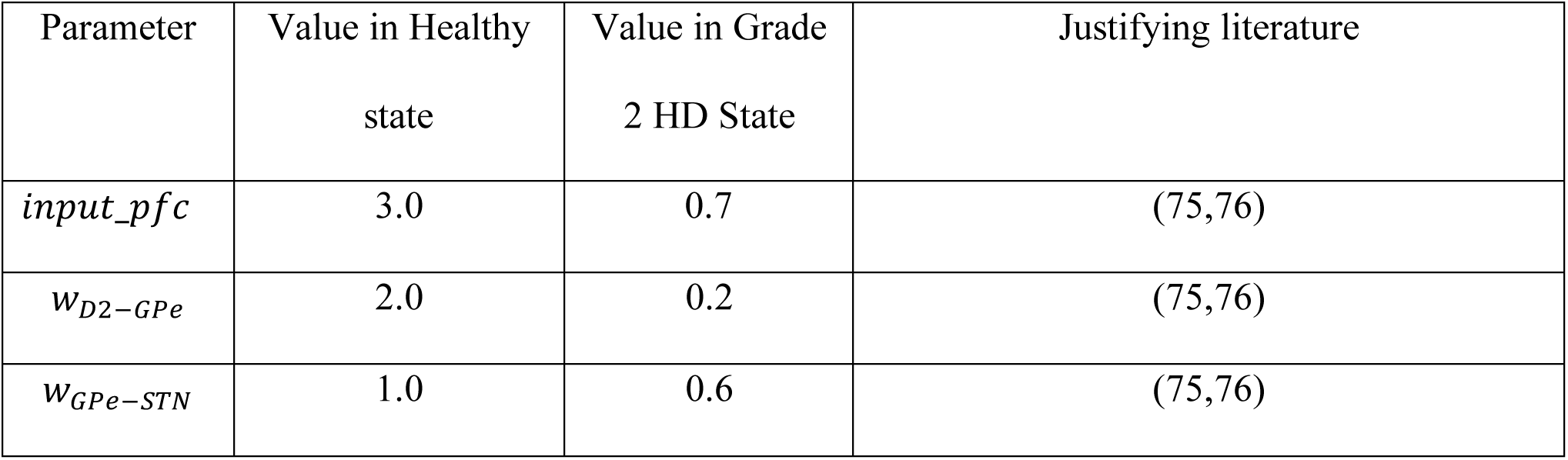
Changes in the parameters of the model that reproduce Huntington disease state.

The cause for the persistent exploratory phase is the positive PMC-BG feedback loop through D1 MSNs, which is not balanced by the D2 MSN pathway. Indeed, an occasional increase in the activity of the PMC2 neural group, which represents a non-rewarded action, excites the corresponding D1 MSN group, and through disinhibition by GPi2 activity, further increases the PMC2 activity (Fig. 9). The reduced connectivity in the D2 MSN pathway makes the STN neural activity the same for choices 1 and 2 (data not shown) and excludes the indirect pathway from the competition between the choices. This leads to occasional choices of the non-rewarded option, and our simulations show that this behavior is robust with respect to growing PFC-PMC and PFC-MSN connections (Fig. 8). Therefore, the lack of balance between direct and indirect pathways in the model of Huntington’s disease causes persistent random switching from rewarded to non-rewarded choice after both initial learning and reversal.

**Figure 9:**
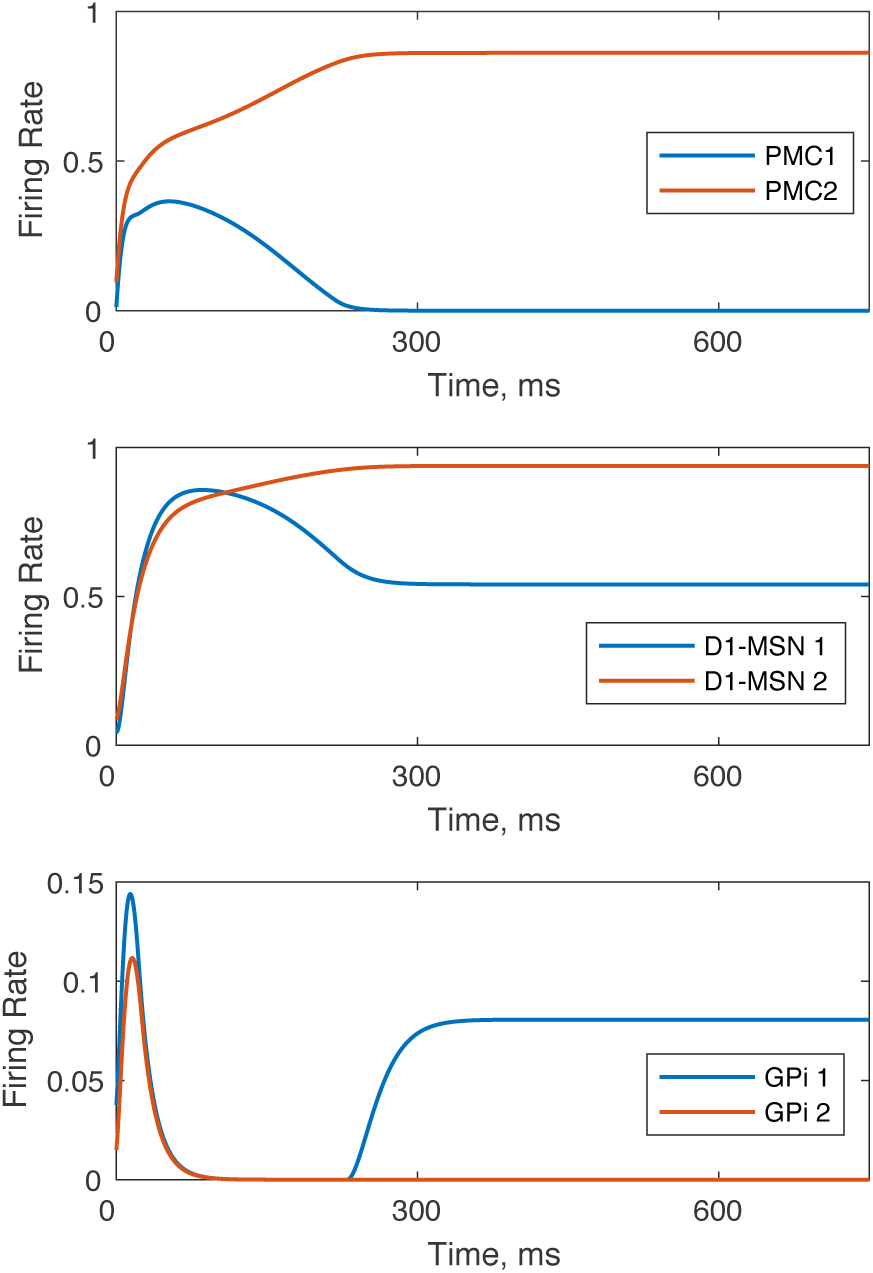
Occasional choice of the nonrewarded option made in the model with Huntington state BG. Within-trial dynamics of PMC, D1 MSN, and GPi neural activity is shown. The greater activity of PMC2 groups signifies that the action 2 is chosen, even though choice 1 is made preferable in the model by potentiating PFC-PMC1, PFC-D1 MSN1 and PFC-D2 MSN2 connections: *W*_*PFC*1–*PMC*1_ = 0.04, *W*_*PFC*1–*D*1*MSN*1_ = 1, *W*_*PFC*1–*D*2*MSN*2_ = 1

In order to model the impact of BG DBS or surgical interventions on performance and learning in HD, we also performed additional simulations of the HD model in which the BG signal to PMC was ablated from trial 100 till the end (Fig. 10). The random switches between the choices cease shortly after, but not at the onset of DBS. The response to DBS is very similar to that in the PD case (Fig. 7). In this period, the PFC-striatal connections no longer exert any influence on the choices, but the PFC-PMC connections are strong enough to lock the choice to the rewarded option. After the reversal on trial 200, however, the changed values of the choices remain unnoticed by the system, the choice remains locked on the now unrewarded option, and the cortical connections supporting this choice keep rising. Therefore, during DBS, or after surgical interventions ablating BG output, behavior improves, but learning is impaired in HD as well as in the PD state.

**Figure 10:**
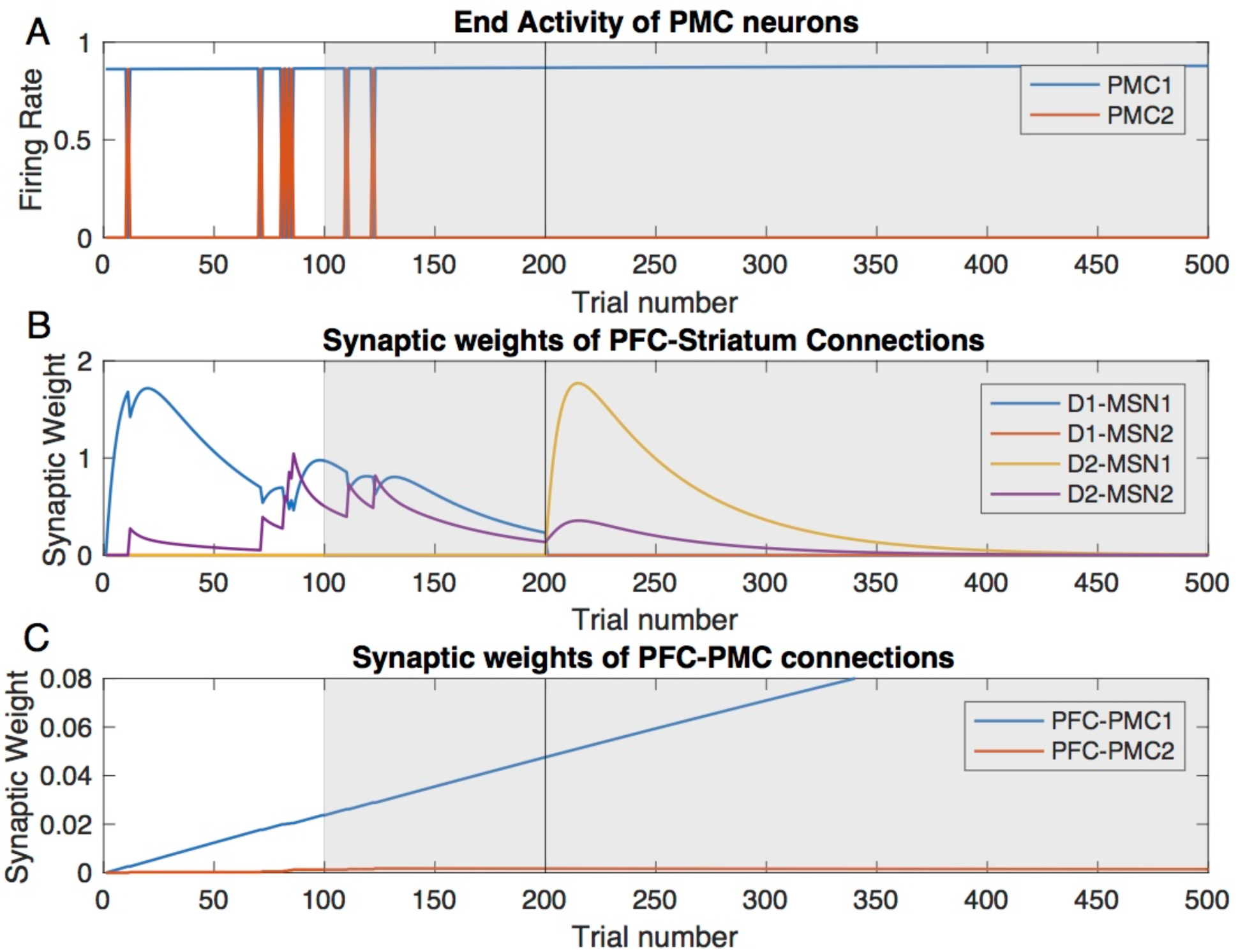
In the HD state, the random switches between choice 1 and 2 cease shortly after, but not at the DBS onset. Trial-by-trial dynamics of the PMC activity and underlying modulation of synaptic weights in the Huntington BG model with simulated DBS starting at trial 100. Same notation as in Fig. 2. (A) The levels of PMC1 and PMC2 activity (choice 1 vs. 2) at the end of each trial (B) Synaptic weights of the PFC to striatum connections reflect rewarded choices. (C) Synaptic weight of the PFC to PMC1 connection keep growing after DBS onset, and during reversal.

## Discussion

Our model implements the cortico-BG-thalamo-cortical loop function in a standard 2-choice instrumental conditioning task. We have shown that potentiation of corticostriatal synapses enables learning of rewarded options. However, later these synapses become redundant as direct connections between prefrontal and premotor cortices (PFC-PMC) potentiate by Hebbian learning. The model shows that disease-related imbalances of the direct and indirect pathways in the BG impairs learning and suggests that these imbalances may also impede choices that have been learned previously, in spite of BG redundancy for those choices.

Our model of the parkinsonian state reproduces several major behavioral and electrophysiological features documented experimentally: First, initial learning is much slower, but reversal takes about as many trials in the mild PD state as it does in the healthy state (∼100 trails). As initial learning is associated with an unpredicted reward (positive RPE) and reversal with reward omission (negative RPE), which is similar to punishment, this is consistent with experimental findings, in which reward, but not punishment learning is impeded in PD patients (47,48). Second, the overall PMC activity is diminished in the PD state, consistent with PD studies (49). Further, the model predicts that this activity is lowest at the beginning of the initial learning and reversal due to aberrant engagement of the indirect pathway, which can be displayed as stronger bradykinesia. Third, the model shows robust oscillations in the activity of the cortico-BG-thalamo-cortical loop in the PD state. The oscillations are generated by a negative feedback branch of the loop through the indirect pathway as suggested before (50,51). The frequency of these oscillations is about 5 Hz, which is in the theta band. An increase in the EEG theta band is a marker of PD-related cognitive decline (52,53). Our simulations show that the oscillations cause multiple choice errors and, consequently, impede task performance and learning.

In the HD state, our model displays persistent randomly occurring choices of the unrewarded option, especially frequent after the reversal. This would register as impaired learning in behavioral tests, which is consistent with experimental results for cognitive (54,55) and motor tasks (56,57) in HD patients in the early stages of the disease. Furthermore, the model suggests that performance for previously learned tasks is also affected.

Therefore, our model reproduces impairments of the previously learned actions documented in BG-affecting diseases like PD and HD as well as after certain BG lesions (8,32,58). However, surgical and deep brain stimulation (DBS) interventions in PD and HD patients do not impair, but rather restore motor function (33–35,59). This raises the question: how can these two lines of evidence therefore be reconciled?

Learning in the model consists of two phases: BG-based and cortex-based. In a faster BG-based phase, the connections from PFC to MSNs are potentiated according to the RPE signal. The BG output inhibits choices with negative RPE and disinhibits those with positive RPE. Once the behavior is learned, the RPE becomes zero, and the PFC-MSN connections decay to zero. The future choices are supported by the slower cortex-based learning phase: The connections from PFC directly to PMC are potentiated based on the Hebbian mechanism. Our simulations show that, even after the cortico-cortical connections increase to the levels ensuring robust choice of the rewarded option in the healthy state, both of the disease models are unable to make robust choices. Thus, behaviors that no longer need the BG are impaired. The model shows that it is an abnormal BG output that impairs the choices. Indeed, the BG output to the PMC does not vanish even when the behavior is learned and the BG no longer receives any RPE signal. In this case, due to the inputs from the PMC, the healthy BG disinhibits the previously learned choice, i.e. it conforms with the PFC-PMC associations. This disinhibitory function is impaired in both PD and HD, as well as after striatal lesions (8,32,58). According to this prediction, disruption of the BG output by GPi lesions or DBS, which was successfully used in PD (33–35) and tested in HD patients (59), would improve performance on previously learned tasks. Indeed, our model of a lesion of BG output demonstrates strengthening of performance on previously learned choices. Therefore our model reconciles how specific GPi lesions or DBS that abolish BG output, restore previously learned behaviors that were lost due to disrupted BG function, however this comes at the expense of decreased cognitive flexibility.

Altogether, we have modeled the function of the cortico-BG-thalamo-cortical loop in a 2 choice instrumental conditioning task and shown the mechanism by which this function is disrupted in HD and PD conditions. Further, we have shown how DBS or GPi lesions restore previously learned choices, but completely disrupts learning of new behavior. Our results reconcile the apparent contradiction between the critical involvement of the BG in execution of previously learned actions and yet no impairment of these actions after BG output is ablated by lesions or DBS.

## Materials and Methods

We adopt rate model formalism extensively used to reproduce activity and function of numerous brain structures (60). In particular, we follow a validated model of motor control (38) and modify it for action selection.

### Structure of the basal ganglia

Fig. 1 presents a schematic diagram of nuclei and connections within the BG and their connections with cortices. The cortico-BG-thalamo-cortical loop is separated into channels selective for each of the two actions of the model (see below). First, the striatum, the primary input structure of the BG, receives excitatory inputs from the prefrontal cortex (PFC) and premotor cortex (PMC) in the cerebrum as well as the thalamus. From the striatum, two competing pathways are activated: a direct pathway (striatum-SNr/GPi) and an indirect pathway (striatum-GPe-STN-SNr/GPi). These two pathways converge at the BG output nuclei, the SNr and GPi, and serve to modulate their activites. In the model SNr and GPi activity are treated as one unit. SNr/GPi activity inhibits a corresponding neural group in the thalamus and PMC and blocks the corresponding action. In the model thalamus and PMC activity is treated as a single unit (PMC/Thal). To execute the action, SNr/GPi activity must decrease and disinhibit the PMC/Thal neurons. In addition, DA neurons in the SNc signal a reward prediction error (RPE), which change synaptic weights of PFC-striatum connections via DA-dependent long-term synaptic potentiation (LTP) and long-term synaptic depression (LTD) to allow for reward-based learning.

### Behavioral task

Our model implements a standard design for intertemporal choice tasks (32). The circuitry shown in Fig. 1 is built to reproduce selection between two actions, one of which is rewarded. A typical task is to learn that, for instance, action 1 is rewarded if a conditioning stimulus (CS) is presented. Then, this task is “reversed”: after learning this contingency, the reward following the same CS is shifted to action 2. Thus, the cortico-BG-thalamo-cortical loop has 2 channels: for choice 1 and 2, except for the PFC that represents the CS and the SNc that represents the unexpected reward. Activation of neural groups 1 and 2 in the PMC/thalamus correspond to execution of action 1 and 2 respectively. Thus, in the model, an action is considered selected if the activity level of the corresponding PMC neural group at the end of a simulated trial is higher than that of the other group. The behavioral readout is if the stimulus-reward contingencies can be learned, and how many trials learning takes.

### Firing rate equations

The activity of every neuron (except the dopaminergic neurons in the SNc) is governed by the following differential equation (38):

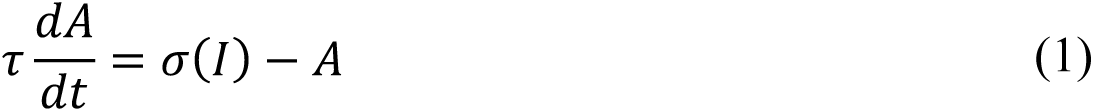

where A is the instantaneous activity level of the neuron. Here, *τ* is a time constant taken to equal 15 msec based on previous models and experimental studies (61). *I* is the synaptic input to the neuron. The expressions for synaptic input to each neuron group, and the formula are compiled in Table 1. *σ*(*I*) is a normalized response function defined as:

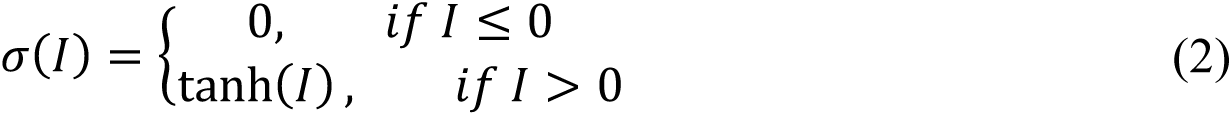

**Table 1:** Synaptic inputs

| Neuron | Formula for Synaptic Input |
| --- | --- |
| PFC | $I_{PFC} = input\_pfc$ |
| D1 MSN | $I_{D1\ MSN_m} = w_{PFC-D1}PFC + w_{PMC-D1}PMC_m$ |
| D2 MSN | $I_{D2\ MSN_m} = w_{PFC-D2}PFC + w_{PMC-D2}PMC_m$ |
| GPe | $I_{GPe_m} = dr_{GPe} - w_{D2-GPe}D2\ MSN_m$ |
| STN | $I_{STN_m} = dr_{STN} - w_{GPe-STN}GPe_m$ |
| GPi | $I_{GPi_m} = dr_{GPi} - w_{D1-GPi}D1\ MSN_m + w_{STN-GPi}STN_m$ |
| PMC | $I_{PMC_m} = dr_{PMC} + w_{PFC-PMC_m}PFC - w_{GPi-PMC}GPi_m - w_{PMC_n-PMC_m}PMC_m$ |

We have adapted the following notation: *X_m_* to denote the activity (firing rate) of neural group X in the pathway for the m^th^ action. Since our model contains only two actions, the only possible values for m are 1 and 2. The index *n* in the formula for *X_m_* refers to the other of the two channels, e.g. 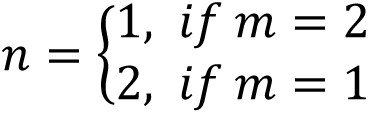. Further, *w_X_Y_* denotes the synaptic weight (strength of connection) from group X to group Y and *dr_X_* denotes a tonic drive to group X. Many of these weights are assumed constant throughout our trials, but several of them are plastic as described below.

### Synaptic plasticity

The synaptic weights from PFC to PMC neurons and from PFC to MSNs are plastic, which means that they change depending on the activity of these nuclei and behavioral outcome (reward received) respectively (40,41,39). In simulations, the synaptic weights are updated at the beginning of every trial depending on the behavior of the model in previous trials. Before we discuss the specific mechanisms by which we updated these plastic synaptic weights, we will first discuss how we calculated the activity of the dopaminergic neurons in the SNc, which essentially mediate reward-based learning.

The activity of the SNc neurons is associated with a reward prediction error (RPE) (62). Following previous models (e.g. (38)), we assume that the activity of the SNc neural group reflects the difference between the expected reward and the actual reward:

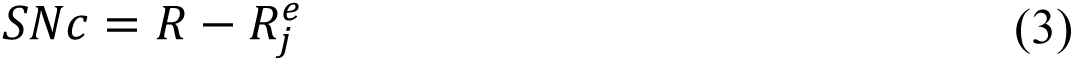

where *R* is the actual reward given based on the action selected, and 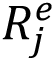 is the expected reward at the j^th^ trial. The expected reward on the first trial, 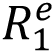, is equal to 0 and is then subsequently updated according to the following scheme:

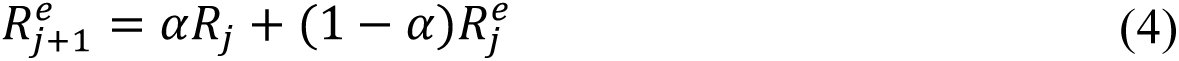

where *α* is a constant (set equal to 0.15) and *R_j_* denotes the actual reward received by the model on the j^th^ trial.

The actual reward received in simulations, *R*, is determined by the following:

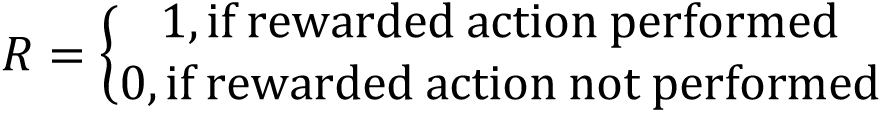

where we determined which action is selected by comparing the activities of the PMC neurons at the end of each trial as described above.

Altogether, after each trial, the PFC-striatal synaptic connections are updated according to the following rules:

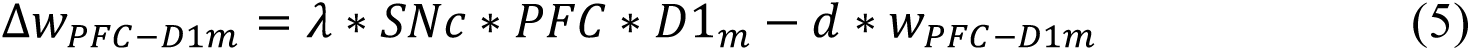

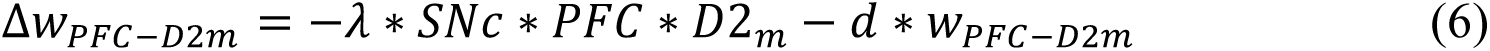

where *λ* is a learning rate constant and *d* is the decay rate constant. Here, *PFC*,*D*1*_m_*, and *D*2*_m_* denote the activity of the respective neural group at the end of the trial (*m* = 1,2).

Lastly, we describe the mechanism by which we updated the connections between the PFC and PMC neurons. Here, we let *W_PFC–PMCm_* denote the synaptic weight of the connection between the PFC neural group and the m^th^ PMC neural group. After each trial, the synaptic weights are updated according to the following Hebbian Learning Rule:

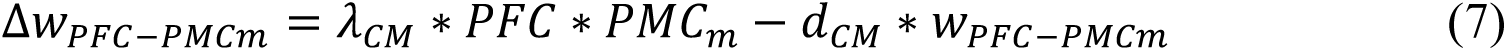

where *λ_CM_* is the learning rate and *d_CM_* is the decay rate of the cortical connections. Here, *PFC* and *PMC_m_* denote the activity of the PFC neurons and m^th^ PMC neuron group at the end of the trial.

Now, we will outline our methodologies for calibrating our three different BG model states: healthy, Parkinsonian, and Huntington’s disease.

### Healthy BG state

We target to reproduce rodent behavior in instrumental conditioning (IC) tasks (32). Thus, an animal will learn contingencies between a conditioning signal and a rewarded action— pressing one of two levers. We reduce the model by (38) and focus our model on the interaction of the thalamocortical and BG networks (Fig. 1) and reproduce the function of the cortico-BG-thalamo-cortical loop in the above two-choice task. The parameter values are shown in Table 1. The values were taken from previous studies (38) with a few minor modifications that allow for both robust instrumental conditioning as well as reversal learning.

### Parkinsonian BG state

To create disease models from our healthy BG model, we reviewed physiological data. The neuropathology of Parkinson’s Disease (PD) is incredibly well-understood: it begins with the destruction of the dopaminergic neurons in the SNc (63,64). Further, the disease is accompanied by a decreased firing rate of the D1 MSNs (65,66), GPe (67–69), and PMC (70) as well as increased firing rates in the D2 MSNs (65,66), STN (71,72), and GPi (73,67,74). We induced an in silico mild Parkinsonian state in our model by suppressing SNc output by 70% and changing synaptic weights along with tonic drives (49,64) as outlined in Table 2.

### Huntington’s BG state

The pathology of Huntington’s Disease (HD) is less well-understood; however, it is clear that there is a progression of the disease from chorea (involuntary, jerky movement) at its onset to akinesia (loss of the power of voluntary movement) at its conclusion (75). We modeled the chorea phase (Grade 2 HD) by weakening the D2 MSN-GPe connection by 90%, weakening the GPe-STN connection by 40%, and decreasing the PFC input to account for destruction of the PFC (75,76). These percentages are gathered from the physiological observations of Reiner et al. (75). The resulting parameters are shown in Table 3.

### Numerical Simulations

Our model was coded in MATLAB. We considered a trial to last 750 msec, and at the end we register the activity of each neuron in the circuit. We chose to cutoff trials at this point because it was sufficient to guarantee that the neural activity converges to a steady state. An exception is a case when neural activity does not approach a steady state and remains oscillatory, which we also found in this study. We update strengths for the plastic synapses after each trial. Finally, we reset the initial activity of the neurons to be at randomized levels at the beginning of each subsequent trial. We ran simulations consisting of 500 such trials.

